# Diversity of Culturable Endophytic bacteria from Wild and Cultivated Rice showed potential Plant Growth Promoting activities

**DOI:** 10.1101/310797

**Authors:** Madhusmita Borah, Saurav Das, Himangshu Baruah, Robin C. Boro, Madhumita Barooah

**Affiliations:** Department of Agricultural Biotechnology, Assam Agricultural University, Jorhat, Assam; Department of Agricultural Biotechnology, Assam Agricultural University, Jorhat, Assam. Email Id

**Keywords:** Endophytes, phytohormone production, mineral solubilization, siderophore, biocontrol, pot culture

## Abstract

In this paper, we report the endophytic microbial diversity of cultivated and wild *Oryza sativa* plants including their functional traits related to multiple traits that promote plant growth and development. Around 255 bacteria were isolated out of which 70 isolates were selected for further studies based on their morphological differences. The isolates were characterized both at biochemical and at the molecular level by 16s rRNA gene sequencing. Based on 16S rRNA gene sequencing the isolates were categorized into three major phyla, viz, Firmicutes (57.1 %), Actinobacteria (20.0 %) and Proteobacteria (22.8 %). Firmicutes was the dominant group of bacteria of which the most abundant genus was *Bacillus*. The isolates were further screened *in vitro* for plant growth promoting activities which revealed a hitherto unreported endophytic bacterial isolate, *Microbacteriaceae bacterium* RS01 11 as the highest secretor of a phytohormone, IAA (28.39 ± 1.39 μg/ml) and GA (67.23 ± 1.83 μg/ml). *Bacillus subtilis* RHS 01 displayed highest phosphate solubilizing activity (81.70 ± 1.98 μg/ml) while, *Microbacterium testaceum* MK LS01, and *Microbacterium trichothecenolyticum* MI03 L05 exhibited highest potassium solubilizing activity (53.42±0.75μg/ml) and zinc solubilizing efficiency (157.50%) respectively. *Bacillus barbaricus* LP20 05 produced highest siderophore units (64.8 %). Potential plant growth promoting isolated were tested *in vivo* in pot culture under greenhouse conditions. A consortium consisting of *Microbacteriaceae bacterium* RS01 11, *Bacillus testaceum* MK LS01 and *Bacillus subtilis* RHS promoted plant growth and increased the yield 3.4 fold in rice when compared to control T0 when tested in pot culture and reduce application rates of chemical fertilizer to half the recommended dose. Our study confirms the potentiality of the rice endophytes isolated as good plant growth promoter and effective biofertilizer.

## INTRODUCTION

In a natural ecosystem, all the healthy and asymptomatic plants host a diverse group of the microbial community including bacteria, fungi, viruses and protista collectively, known as plant microbiota (Hiruma et al., 2016). Among the plant-associated microorganisms, endophytes are the bacterial and fungal population colonizing within a plant tissue for a part of its life cycle without showing any apparent pathogenesis (Tan and Zou, 2001). Culture-dependent and independent community profiling revealed their active association virtually with all the tissues of a host plant, including the intercellular spaces of the cell walls, vascular bundles, and in reproductive organs of plants, e.g. flowers, fruits, and seeds. Their association was even logged from aseptically regenerated tissues of micro-propagated plants (Dias et al., 2009). Environmental parameter including soil nutrients and different abiotic stresses influence the diversification of the endophytic entity in a plant may play a significant role in the natural fitness in particular environment (Bulgarelli et al., 2013; Kogel et al., 2006). In this mutualistic relationship, the plant provides primary nutritive components and a protective niche for the endophytic organisms whereas, the endophytes produce useful metabolites and systemic signals (Rosenblueth and Martínez-Romero, 2006; Strobel, 2003). Endophytic bacteria like *Bacillus, Enterobacter, Klebsiella, Pseudomonas, Burkholderia, Pantoea, Agrobacterium, etc*. have been isolated from diverse plant species including maize, potato, tomato, sugarcane, and cucumber (Bacon and Hinton, 2007).

Although the endophytic relationship was documented long ago by Perotti, (1926), many aspects of this mutualistic relationship are poorly understood including the molecular mechanisms underlying such association and the selective association of a particular group of endophytes(Xia et al., 2015). Most of the reports on endophytic colonization in plants have focused on plant and root endophytic association (Lundberg et al., 2013; Romero et al., 2014). Plant-microbe association has been studied for many decades for sustainable agricultural practices. Endophytes are known for their ability to promote plant growth either directly or indirectly through several metabolic activities including facilitating the acquisition of mineral resources like phosphorus, potassium, zinc, and iron or by regulating the phytohormone production including auxin, gibberellin, and cytokinin (Glick, 2014; Rosenblueth and Martínez-Romero, 2006). Indirectly, they can stimulate host growth by antagonistic activity or by inducing systemic resistance against different phytopathogens (Arnold, 2007; Pillay and Nowak, 1997). A particular endophyte can affect the plant growth and development using one or more of these mechanisms.

Of the nearly 3,00,000 plant species that exist on the earth, each individual plant is host to one or more endophytes but only a few of these plants have ever been completely studied relative to their endophytic biology (Strobel et al., 2004). Thus, the probability of isolation of novel and beneficial endophytic microorganisms from the diverse flora is considerably high. Plants growing in areas of biodiversity hotspot may be host to endophytes hitherto unreported. Assam is located within the Indo-Burma biodiversity hotspot and a secondary center of *Oryza sativa* with more than 4000 accessions of germplasm. Along with the cultivated rice varieties, Assam harbors a significantly high number of wild accessions mainly belonging to *Oryza rufipogon*. Thus, the wild rice together with cultivated ones can be a potential host to the different endophytic community with eco-physiological characteristics for adaption to different biotic and abiotic stresses. Exploration of endophyte-plant interaction can help to devise a low-input sustainable agricultural application for different crops in various farming conditions. Thus, in this paper, we report the diverse endophytic community of rice through culture-dependent profiling and characterization of the potent endophytes for their plant growth promoting activity and further present results of their influence to promote crop growth and yield.

## Materials and Methods

### Isolation and Characterization

Locally cultivated rice varieties *viz*., Kola Joha, Miatong, Barjahi, and Gitesh were collected at booting stage from Jorhat (26°43′03.8″N 94°11′40.2″E), Tinsukia (27°20′34.5″N 95°42′33.2″E) and Lakhimpur (26°57′38.5″N 93°51′53.5″″E) districts of Assam. In addition to cultivated plants (*Oryza sativa*) different morphotypes of wild rice *O. rufipogon*, locally known as “*Uri-DhoF*” were collected to assess the endophytic diversity of prokaryotic microorganisms by the culture-dependent approach. Healthy and disease free paddy samples were selected, uprooted from rice fields and immediately transported to the laboratory in ice boxes. The plant samples were thoroughly cleaned with running water to remove the attached debris. After that, leaves, stems, and roots were separated and cut into thin sections of 2-3 cm long and washed thoroughly with double distilled water. The samples were rinsed in 70% ethanol, sterilized with 0.1% HgCl_2_ and further washed with sterile distilled water for several times to remove the surface sterilizing agents (Gagné et al., 1987). One gram of the samples were homogenized in 10 ml of distilled water to prepare a stock solution of tissue homogenate. The appropriate diluted sample was inoculated in Tryptic Soya Agar (TSA) plates and incubated at 30° C for 48 hrs and pure cultures were isolated by streak plate method. The bacterial isolates were characterized both morphologically and biochemically through various tests (gram staining, starch hydrolysis, casein hydrolysis, catalase reaction, citrate and malate utilization, nitrate reduction, H2S production and gelatin liquefaction) according to the Bergey’s Manual of Determinative Bacteriology (Krieg, 2015).

### Molecular Characterization

Genomic DNA was extracted from bacteria as per standard phenol-chloroform method. The 1500 bp region of the 16S rRNA gene was amplified from the extracted genomic DNA using the universal forward primer 5′-AGAGTTTGATCCTGGCTC −3′ and reverse primer 5′-AAGGAGGTGATCCAGCCG-3′. The PCR products thus obtained were sequenced. The forward and reverse sequences obtained were assembled using the Codon Code Aligner software. Nucleotide sequence identities were determined using the BLAST tool from the National Center for Biotechnology Information (NCBI). Partial sequence data for the 16S rRNA genes have been deposited in the Gen Bank nucleotide sequence data libraries and Gene Bank accession numbers have been provided to these sequences. After aligning the sequence of the 16S rRNA region, a phylogenetic tree was constructed using MEGA 6.0 based on neighbor-joining method for the analysis of evolutionary relatedness and the evolutionary distances were computed using the Kimura 2-parameter method (Kimura, 1980).

### Determination of Species Diversity

Diversity and the relative species abundance of the endophytic isolates identified in this study were calculated using the Shannon diversity index and Simpson diversity index. The PAST software program was adopted to measure the ecological diversity indices and generate the rarefaction curves to evaluate the overall richness (Ryan et al., 2001).

### In vitro Plant Growth Promoting Traits Phytohormone Production

The endophytic bacterial isolates were screened for *in-vitro* phytohormone production mainly Indole Acetic Acid (IAA) and Gibberellic Acid (GA) Quantitative estimation of IAA and GA was determined by following the method described earlier (Patten and Glick, 2002; Vikram et al., 2007).

### Mineral Solubilization

The isolates were checked for their different mineral solubilizing activities including phosphate, potassium and zinc solubilization by following the method described earlier (Hu et al., 2006; Ramesh et al., 2014).

### Siderophore Production

Bacterial isolates were assayed for siderophore production as described by Schwyn and Neilands (1987) (Schwyn and Neilands, 1987). Quantitative estimation of siderophores was done by CAS shuttle assay (Payne, 1994).

### Efficacy of bio-inoculum on plant growth promotion under greenhouse conditions Plant material

The rice cultivar used for the pot culture study was Dichang, which is a short duration variety. Dichang can be grown in Sali (June/ July to November/ December) and Boro (November/ December to May/June) season; however, we opted for the Boro season to carry out our experiment.

### Inoculum Preparation

Identified bacterial strains showing high phytostimulent activity viz. *Microbacteriaceae bacterium* RS01 11, *Bacillus subtilis* RHS01 and *Microbacterium testaceum* MK LS01 were analyzed for evaluating the efficacy of the strains in an *in-vivo* pot culture experiment under greenhouse condition. Culture inoculum was prepared by mixing equal quantities of each culture just before application. Prior to preparation of consortia, compatibility of isolates was checked according to Fukui (1994).

### Experimental design

The experiment was arranged in a complete randomized design (CRD) with five replications per treatment. The test was performed with six different treatments with compositional differences in bio inoculum, organic compost (vermicompost) and inorganic fertilizers. A control set of pot was maintained without any treatment to replicate the controlled environment. Treatments designed were T0: Soil (Untreated); T1: Soil + Chemical fertilizer (NPK); T2: Soil + Vermicompost; T3: Soil + Bioinoculum; T4: Soil + Vermicompost + Bioinoculum; T5: Soil + ½ NPK + ½ Vermicompost; T6: Soil + ½ NPK + ½ Vermicompost + ½ Bioinoculum.

### Pot preparation

Soil from rice farming fields was collected, air-dried and sieved. Chemical fertilizer Nitrogen, Phosphorus, and Potassium (NPK) were used in a ratio of 40:20:20 kg/hectare. Vermicompost was used in a ration of 300gm/10kg of soil. In bioinoculum treatment, each pot received 20 ml of bacterial inoculum.

### Seed sowing and harvesting of the plants

Rice seeds were soaked in sterilized water in a Petri dish for 24 hours. The water was drained off and the seeds were kept in a closed Petri dish in warm conditions for 2 days. Four pre-germinated seeds were allowed to grow in each pot in the greenhouse. The plants were watered twice a day to maintain optimum soil moisture regime and kept under greenhouse condition with ambient temperature and air humidity. The plant was regularly monitored till harvest (150 days) for gradual growth promotion. Parameters selected for assessing the growth of the plant were plant height, number of tillers, number of leaves per tiller, length of the flag leaf, number of panicles per tiller, the total number of seeds per plant, weight of 100 grains, weight of dry biomass and yield per plant. Plant growth parameters were measured from 30 days till harvest in an interval of every 15 days

### Statistical analysis

Data from the quantitative analysis of plant growth promoting traits and pot culture experiment were analyzed by one-way analysis of variance (ANOVA). Statistical analysis was performed by using SPSS software (version 18). Significant differences between means were compared using least significant differences test (LSD) at 5% (*p* ≤ 0.05) probability level.

## Result

### Isolation and Characterization

A total of 255 bacterial endophytes were isolated from leaf, stem and root section of both cultivated and wild rice plant. About 70 different isolates were selected on the basis of their varying morphological characteristics. Bacterial endophytes were characterized biochemically, 53 isolates were found to be gram-positive and 17 gram-negative. Thirty-three isolates were positive for starch hydrolysis (amylase producers), 15 isolates were found positive for casein hydrolysis (protease producers), 41 isolates were catalase producers, 14 citrate utilizers, 35 malate utilizers, 22 nitrate reducers, 33 H2S producers and 46 gelatinase producers (**Table 1**).

**Table 1:** Morphological and biochemical characterization of the isolates.

| Sl N o. | Sample Code | Cell Shape | Gram reaction | Starch Hydrolysis | Caesin Hydrolysis | Catalase Reaction | Citrate Utilization | Malate Utilization | Nitrate Reduction | H<sub>2</sub>S Production | Gelatin liquefaction | Cellulase Production | Pectinase Production |
| --- | --- | --- | --- | --- | --- | --- | --- | --- | --- | --- | --- | --- | --- |
| 1. | MI3 L05 | rod | + | - | - | - | - | - | + | - | - | - | - |
| 2. | RR01 3D | rod | + | + | - | + | - | + | - | - | + | - | + |
| 3. | RR01 04 | rod | + | + | + | + | - | + | - | + | + | - | - |
| 4. | RS01 05 | rod | + | - | - | + | - | - | - | - | + | + | - |
| 5. | RS01 11 | rod | + | + | - | + | - | + | - | + | - | + | - |
| 6. | RLS 04 | rod | + | + | - | - | + | - | + | + | + | - | + |
| 7. | RHS 06 | rod | + | - | - | + | + | - | + | - | + | - | - |
| 8. | RHS 01 | rod | + | + | - | + | + | + | + | + | - | - | - |
| 9. | RHS 11 | rod | + | - | + | - | - | + | + | + | + | + | + |
| 10. | RLS 12 | rod | + | + | + | + | + | + | + | + | + | + | - |
| 11. | LP10 S10 | rod | + | + | + | + | + | + | + | + | + | + | - |
| 12. | LP10 S01 | rod | - | - | - | + | - | + | - | - | - | - | - |
| 13. | LP21 S03 | rod | + | - | - | - | - | - | + | - | - | + | - |
| 14. | LP31 R13 | round | - | - | - | - | + | - | + | - | - | - | - |
| 15. | LP10 L02 | *rod* | - | - | - | - | - | - | + | + | + | + | - |
| 16. | RHS 02 | *rod* | - | - | + | - | - | - | + | + | + | - | - |
| 17. | LP21 R02 | rod | + | - | - | - | - | - | + | + | - | - | - |
| 18. | LP10<br>S06 | rod | + | + | + | + | - | + | - | - | + | - | - |
| 19. | M12<br>LS01 | rod | + | + | - | - | - | - | - | - | + | - | - |
| 20. | LP01<br>R08 | rod | + | + | - | + | - | - | - | - | - | - | - |
| 21. | MK2L02 | rod | + | - | - | - | - | - | + | - | - | - | - |
| 22. | LP21S02 | rod | + | - | - | - | - | - | + | - | - | - | - |
| 23. | LP01L06 | rod | + | + |  | + | - | - | - | - | + | - | - |
| 24. | LP35L05 | rod | + | - | + | + | - | + | - | - | + | - | - |
| 25. | LP31L19 | *rod* | - | - | - | - | - | - | - | - | - | - | - |
| 26. | LP01S02 | round | + | - | - | - | - | - | - | - | + | - | - |
| 27. | LP10L02 | *rod* | - | - | + | - | - | - | - | + | + | - | - |
| 28. | RHS07 | round | - | - | + | - | - | - | - | + | - | - | - |
| 29. | LP31<br>R11 | rod | + | + | - | + | + | + | - | - | + | - | - |
| 30. | LP31<br>L04 | rod | + | + | + | + | - | + | - | - | + | - | - |
| 31. | LP35<br>L05 | rod | + | - | + | + | - | + | - | - | + | - | - |
| 32. | RHD 11 | *rod* | - | - | - | - | - | - | - | + | + | - | - |
| 33. | RR01 3C | rod | - | - | - | + | - | - | + | - | - | - | - |
| 34. | RS01 04 | rod | - | - | - | - | + | - | - | - | - | - | - |
| 35. | RHM<br>104 | rod | + | + | - | + | - | + | - | + | + | - | + |
| 36. | RHM<br>105 | rod | + | + | - | + | - | + | - | + | + | - | - |
| 37. | RHM 109 | rod | + | + | - | + | - | + | - | + | - | + | - |
| 38. | RHM 111 | rod | + | + | + | + | - | + | - | + | + | - | - |
| 39. | RHM 113 | rod | + | + | - | + | - | + | - | + | + | - | + |
| 40. | RHM 120 | rod | + | + | - | + | - | + | - | + | + | + | - |
| 41. | RHM 122 | rod | + | + | - | + | - | + | - | + |  | + | - |
| 42. | RHM 123 | rod | + | + | - | + | - | + | - | + | + | + | - |
| 43. | RSC 04 | rod | + | + | - | + | - | + | - | + | + | + | + |
| 44. | RSC 05 | rod | + | + | - | + | - | + | - | + | + | - | - |
| 45. | RLC 11 | rod | + | + | - | + | - | + | - | + | - | - | - |
| 46. | RHC 13 | rod | + | + | - | + | - | + | - | + | + | + | + |
| 47. | RRC 23 | rod | + | - | - |  | - |  | - | - | - | - | - |
| 48. | RRC 24 | rod | + | + | - | + | - | + | - | + | + | - | - |
| 49. | RRC 26 | rod | + | - | - | - | - | - | - | - | - | - | - |
| 50. | MK LS01 | *rod* | + |  | - | - | - | - | - | - | - | + | - |
| 51. | RNS 01 | *rod* | + | + | - | + | - | + | - | + | + | - | - |
| 52. | RNS 02 | *rod* | - | - | - | + | + | - | - | - | + | - | - |
| 53. | RNS 03 | *rod* | + | + | - | + | - | + | - | - | + | - | - |
| 54. | LP02 01 | *rod* | + | - | - | + | - | + | - | - | + | - | - |
| 55. | LP05 05 | rod | - | - | - | - | - | - | + | - | - | - | - |
| 56. | LP05 06 | rod | - | - | - | - | - | - | + | - | - | - | - |
| 57. | LP05 07 | round | + | - | - | - | - | - | + | - | - | - | - |
| 58. | LP05 08 | round | + | - | - | - | - | - | + | - | - | - | - |
| 59. | LP12 05 | rod | + | - | - | - | - | - | + | - | - | - | - |
| 60. | LP12 09 | rod | + | - | - | - | - | - | + | - | - | - | - |
| 61. | LP12 10 | rod | + | - | - | - | - | - | - | - | - | - | - |
| 62. | LP12 11 | rod | - | - | - | + | + | - | - | + | + | - | - |
| 63. | LP20 01 | rod | + | + | - | + | - | + | - | + | + | - | - |
| 64. | LP20 03 | rod | + | + | + | + | + | + | - | + | + | - | - |
| 65. | LP20 04 | *rod* | + | + | + | + | + | + | - | + | + | - | - |
| 66. | LP20 05 | *rod* | + | + | - | + | - | + | - | + | + | - | - |
| 67. | LP20 06 | *rod* | - | - | - | - | - | - | - | - | - | - | - |
| 68. | RNS 04 | rod | - | - | + | - | - | - | + | + | + | - | - |
| 69. | RNS 05 | rod | - | - | - | + | - | - | - | - | + | - | - |
| 70. | LP31<br>L03 | rod | + | + | - | + | - | - | - | - | + | - | - |

Isolates were further characterized at the molecular level using 16S rRNA gene sequence data, 70 different bacterial endophytic strains were identified and submitted to GenBank Database and GenBank accession numbers obtained (**Table 2**). A phylogenetic tree was constructed using the 16S rRNA sequences (**Fig 1**) and the evolutionary history was inferred using the Neighbor-Joining method in MEGA6. The bacteria isolated in this study belonged to 3 major phyla, *viz*., Firmicutes (57.1 %), Actinobacteria (20.0 %) and Proteobacteria (22.8 %). Isolates from cultivated rice and wild rice were grouped together as they share the same phylogenetic origin. Gram-positive bacteria and gram-negative formed two major independent clusters, Cluster I and Cluster II respectively. The Cluster I included 2 plylums *viz*. Firmicutes and Actinobacteria while Cluster 2 comprised of phylum, Proteobacteria. Within the Firmicutes the major clade belonged to the class Bacilli mainly encompassing the genus *Bacillus* (92.5%). Actinobacteria, the second clade of gram-positive bacteria encompassed members of the class Actinobacteria under which genus Microbacterium, Microbacteriaceae and Cellulosimicrobium were observed. Analysis of the Proteobacteria revealed the presence of Alphaproteobacteria, Betaproteobacteria and Gammaproteobacteria. Majority of the clades were affiliated with Gammaproteobacteria, mostly by Pseudomonaceae (43.75%). Other clades of some minor groups such as Enterobacteriaceae, Moraxellaceae, Xanthomonadaceae, Burkholderiaceae etc were also observed under this major group.

**Fig. 1:**
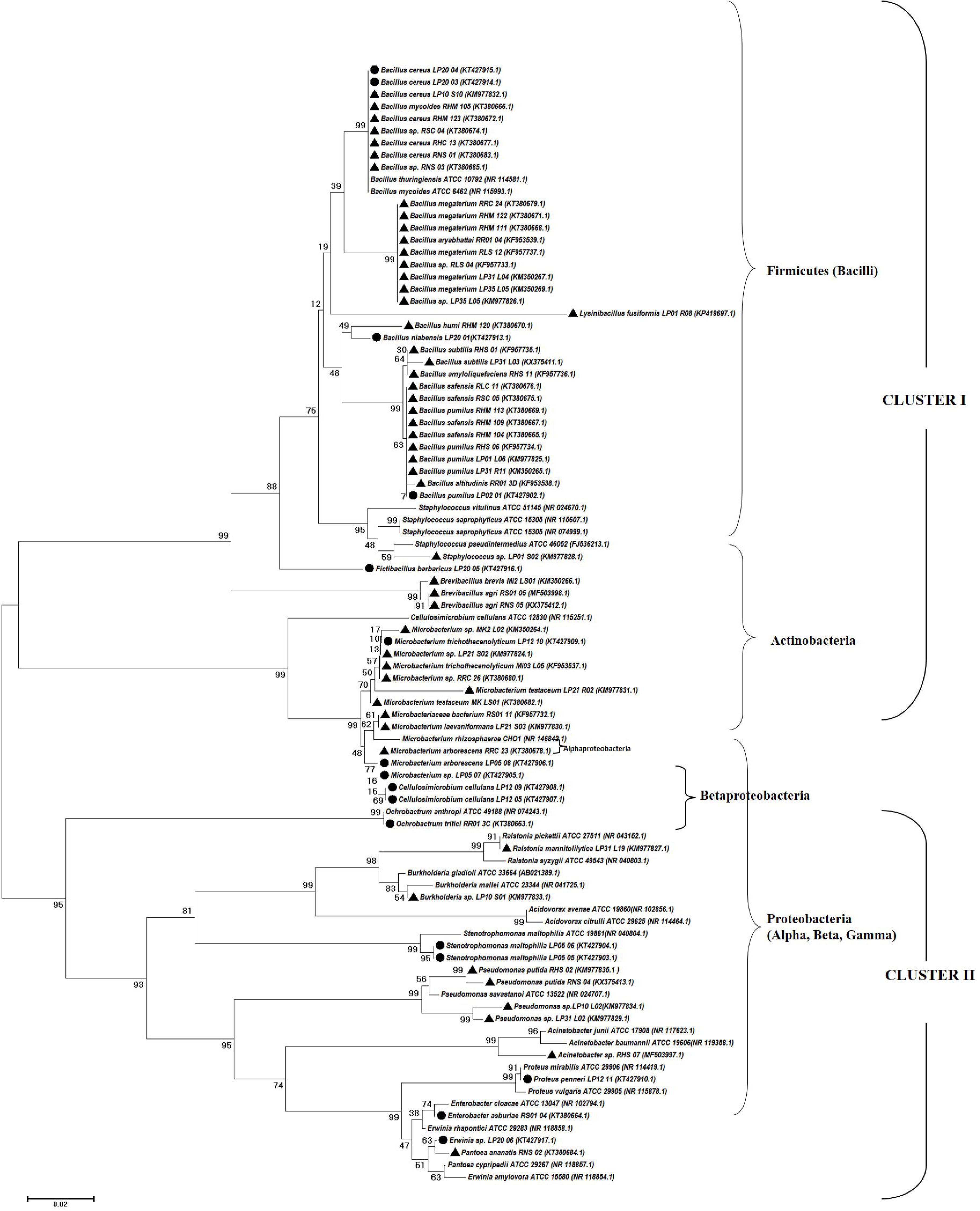
Phylogenetic analysis of 16S rRNA gene sequences of the bacterial isolates along with the reference sequences from NCBI. The analysis was conducted using neighbor-joining method.

**Table 2:** Endophytic bacteria with their isolation source and NCBI accession number.

| Sl. No. | Isolate Code | Isolation Source |  | Organism Match | Family | Phylum | Accession number |
| --- | --- | --- | --- | --- | --- | --- | --- |
|  |  | Type | Plant Part |  |  |  |  |
| 1. | MI03 L05 | Cultivated | Leaf | *Microbacterium trichothecenolyticum* | *Microbacteriaceae* | Actinobacteria | KF953537 |
| 2. | RR01 3D | Cultivated | Root | *Bacillus altitudinis* | Bacillaceae | Firmicutes | KF953538 |
| 3. | RR01 04 | Cultivated | Root | *Bacillus aryabhatai* | Bacillaceae | Firmicutes | KF953539 |
| 4. | RS01 05 | Cultivated | Stem | *Brevibacillus agri* | Bacillaceae | Firmicutes | MF503998 |
| 5. | RS01 11 | Cultivated | Stem | *Microbacteriaceae bacterium* | *Microbacteriaceae* | Actinobacteria | KF957732 |
| 6. | RLS 04 | Cultivated | Leaf | *Bacillus* sp. | Bacillaceae | Firmicutes | KF957733 |
| 7. | RHS 06 | Cultivated | Stem | *Bacillus pumilus* | Bacillaceae | Firmicutes | KF957734 |
| 8. | RHS 01 | Cultivated | Stem | *Bacillus subtilis* | Bacillaceae | Firmicutes | KF957735 |
| 9. | RHS 11 | Cultivated | Stem | *Bacillus amyloliquefaciens* | Bacillaceae | Firmicutes | KF957736 |
| 10. | RLS 12 | Cultivated | Leaf | *Bacillus megaterium* | Bacillaceae | Firmicutes | KF957737 |
| 11. | LP10 S10 | Cultivated | Stem | *Bacillus cereus* | Bacillaceae | Firmicutes | KM977832 |
| 12. | LP10 S01 | Cultivated | Stem | *Burkholderia* sp. | Burkholderiaceae | Proteobacteria | KM977833 |
| 13. | LP21 S03 | Cultivated | Stem | *Microbacterium laevaniformans* | *Microbacteriaceae* | Actinobacteria | KM977830 |
| 14. | LP31 R13 | Cultivated | Root | *Acinetobacter guillourie* | Moraxellaceae | Proteobacteria | KM977837 |
| 15. | LP10 L02 | Cultivated | Leaf | *Pseudomonas* sp. | Pseudomonadaceae | Proteobacteria | KM977829 |
| 16. | RHS 02 | Cultivated | Stem | *Pseudomonas putida* | Pseudomonadaceae | Proteobacteria | KM977835 |
| 17. | LP21 R02 | Cultivated | Root | *Microbacterium testaceum* | *Microbacteriaceae* | Actinobacteria | KM977831 |
| 18. | LP10 S06 | Cultivated | Stem | *Bacillus cereus* | Bacillaceae | Firmicutes | KM350268 |
| 19. | M12 LS01 | Cultivated | Root | *Brevibacillus brevis* | Paenibacillaceae | Firmicutes | KM350266 |
| 20. | LP01 R08 | Cultivated | Root | *Lysenibacillus fusiformis* | Bacillaceae | Firmicutes | KP419697 |
| 21. | MK2 L02 | Cultivated | Leaf | *Microbacterium* sp. | *Microbacteriaceae* | Actinobacteria | KM350264 |
| 22. | LP21 S02 | Cultivated | Stem | *Microbacterium* sp. | *Microbacteriaceae* | Actinobacteria | KM977824 |
| 23. | LP01 L06 | Cultivated | Leaf | *Bacillus pumilus* | Bacillaceae | Firmicutes | KM977825 |
| 24. | LP35 L05 | Cultivated | Leaf | *Bacillus* sp. | Bacillaceae | Firmicutes | KM977826 |
| 25. | LP31 L19 | Cultivated | Leaf | *Ralstonia mannitolilytica* | Pseudomonadaceae | Proteobacteria | KM977827 |
| 26. | LP01 S02 | Cultivated | Stem | *Staphylococcus* sp. | Staphylococcaceae | Firmicutes | KM977828 |
| 27. | LP10 L02 | Cultivated | Leaf | *Pseudomonas* sp. | Pseudomonadaceae | Proteobacteria | KM977834 |
| 28. | RHS 07 | Cultivated | Stem | *Acinetobacter* sp. | Pseudomonadaceae | Proteobacteria | MF503997 |
| 29. | LP31 R11 | Cultivated | Root | *Bacillus pumilus* | Bacillaceae | Firmicutes | KM350265 |
| 30. | LP31 L04 | Cultivated | Leaf | *Bacillus megaterium* | Bacillaceae | Firmicutes | KM350267 |
| 31. | LP35 L05 | Cultivated | Leaf | *Bacillus megaterium* | Bacillaceae | Firmicutes | KM350269 |
| 32. | RHD 11 | Cultivated | Root | *Pseudomonas* sp. | Pseudomonadaceae | Proteobacteria | KP419696 |
| 33. | RHM 104 | Cultivated | Root | *Bacillus safensis* | Bacillaceae | Firmicutes | KT380665 |
| 34. | RHM 105 | Cultivated | Root | *Bacillus mycoides* | Bacillaceae | Firmicutes | KT380666 |
| 35. | RHM 109 | Cultivated | Root | *Bacillus safensis* | Bacillaceae | Firmicutes | KT380667 |
| 36. | RHM 111 | Cultivated | Root | *Bacillus megaterium* | Bacillaceae | Firmicutes | KT380668 |
| 37. | RHM 113 | Cultivated | Root | *Bacillus pumilus* | Bacillaceae | Firmicutes | KT380669 |
| 38. | RHM 120 | Cultivated | Root | *Bacillus humi* | Bacillaceae | Firmicutes | KT380670 |
| 39. | RHM 122 | Cultivated | Stem | *Bacillus megaterium* | Bacillaceae | Firmicutes | KT380671 |
| 40. | RHM 123 | Cultivated | Root | *Bacillus cereus* | Bacillaceae | Firmicutes | KT380672 |
| 41. | RSC 04 | Cultivated | Stem | *Bacillus* sp. | Bacillaceae | Firmicutes | KT380674 |
| 42. | RSC 05 | Cultivated | Stem | *Bacillus safensis* | Bacillaceae | Firmicutes | KT380675 |
| 43. | RLC 11 | Cultivated | Leaf | *Bacillus safensis* | Bacillaceae | Firmicutes | KT380676 |
| 44. | RHC 13 | Cultivated | Root | *Bacillus cereus* | Bacillaceae | Firmicutes | KT380677 |
| 45. | RRC 23 | Cultivated | Root | *Microbacterium arborescens* | *Microbacteriaceae* | Actinobacteria | KT380678 |
| 46. | RRC 24 | Cultivated | Root | *Bacillus megaterium* | Bacillaceae | Firmicutes | KT380679 |
| 47. | RRC 26 | Cultivated | Root | *Microbacterium* sp. | *Microbacteriaceae* | Actinobacteria | KT380680 |
| 48. | MK LS01 | Cultivated | Leaf | *Microbacterium testaceum* | *Microbacteriaceae* | Actinobacteria | KT380682 |
| 49. | RNS 01 | Cultivated | Leaf | *Bacillus cereus* | Bacillaceae | Firmicutes | KT380683 |
| 50. | RNS 02 | Cultivated | Leaf | *Pantoea ananatis* | Enterobacteriaceae | Proteobacteria | KT380684 |
| 51. | RNS 03 | Cultivated | Root | *Bacillus* sp. | Bacillaceae | Firmicutes | KT380685 |
| 52. | LP02 01 | Wild | Leaf | *Bacillus pumilus* | Bacillaceae | Firmicutes | KT427902 |
| 53. | LP05 05 | Wild | Leaf | *Stenotrophomonas maltophilia* | Xanthomonadaceae | Proteobacteria | KT427903 |
| 54. | LP05 06 | Wild | Stem | *Stenotrophomonas maltophilia* | Xanthomonadaceae | Proteobacteria | KT427904 |
| 55. | LP05 07 | Wild | Root | *Microbacterium* sp. | *Microbacteriaceae* | Actinobacteria | KT427905 |
| 56. | LP05 08 | Wild | Root | *Microbacterium arborescens* | *Microbacteriaceae* | Actinobacteria | KT427906 |
| 57. | LP12 05 | Wild | Leaf | *Cellulosimicrobium cellulans* | *Microbacteriaceae* | Actinobacteria | KT427907 |
| 58. | LP12 09 | Wild | Root | *Cellulosimicrobium cellulans* | *Microbacteriaceae* | Actinobacteria | KT427908 |
| 59. | LP12 10 | Wild | Stem | *Microbacterium trichothecenolyticum* | *Microbacteriaceae* | Actinobacteria | KT427909 |
| 60. | LP12 11 | Wild | Root | *Proteus penneri* | Enterobacteriaceae | Proteobacteria | KT427910 |
| 61. | LP20 01 | Wild | Leaf | *Bacillus niabensis* | Bacillaceae | Firmicutes | KT427913 |
| 62. | LP20 03 | Wild | Root | *Bacillus cereus* | Bacillaceae | Firmicutes | KT427914 |
| 63. | LP20 04 | Wild | Stem | *Bacillus cereus* | Bacillaceae | Firmicutes | KT427915 |
| 64. | LP20 05 | Wild | Stem | *Bacillus barbaricus* | Bacillaceae | Firmicutes | KT427916 |
| 65. | LP20 06 | Wild | Root | *Erwinia* sp. | Enterobacteriaceae | Proteobacteria | KT427917 |
| 66. | RR01 3C | Wild | Root | *Ochrobactrum tritici* | Brucellaceae | Proteobacteria | KT380663 |
| 67. | RS01 04 | Wild | Stem | *Enterobacter asburiae* | Enterobacteriaceae | Proteobacteria | KT380664 |
| 68. | RNS 04 | Cultivated | Stem | *Pseudomonas putida* | Pseudomonadaceae | Proteobacteria | KX375413 |
| 69. | RNS 05 | Cultivated | Stem | *Brevibacillus agri* | Paenibacillaceae | Firmicutes | KX375412 |
| 70. | LP31 L03 | Cultivated | Leaf | *Bacillus subtilis* | Bacillaceae | Firmicutes | KX375411 |

**Table 3:** Diversity indices of endophytes isolated from cultivated and rice.

| Cultivated |  |  |  |  | Wild |  |  |  |
| --- | --- | --- | --- | --- | --- | --- | --- | --- |
|  | Taxa | Individual | Shannon | Simpson | Taxa | Individual | Shannon | Simpson |
|  | S | s | H | 1-D | S | l | nH | n |
|  |  |  |  |  |  | S |  | 1-D |
| **Root** | 16 | 20 | 2.718 | 0.930 | 7 | 7 | 1.946 | 0.857 |
| **Stem** | 15 | 18 | 2.659 | 0.925 | 5 | 5 | 1.609 | 0.800 |
| **Leaf** | 12 | 16 | 2.393 | 0.898 | 4 | 4 | 1.386 | 0.750 |

**Table 4:** Growth characteristics of pot culture till harvest.

| Treatment | Shoot Height (cm) | No. of Tillers | No. of Leaves | Length of flag leaf (cm) | No. of panicles per plant | Total no. of seeds per plant | Weight / 100 seeds (gm) | Dry Biomass of the plant (gm) | Yield/plant |
| --- | --- | --- | --- | --- | --- | --- | --- | --- | --- |
| **T0** | 18.00 c | 4.60 d | 3.40 d | 21.74 e | 6.60 e | 178.00 g | 1.69 e | 16.79 f | 3.00 g |
| **T1** | 22.12 b | 8.20 a | 6.40 a | 39.18 a | 12.20 c | 372.20 c | 1.87 c | 42.78 a | 6.97 c |
| **T2** | 21.06 b | 6.40 b c | 5.40 b | 35.70 b | 10.20 cd | 273.20 e | 1.89 c | 31.18 c | 5.18 e |
| **T3** | 22.24 b | 5.60 c | 4.40 c | 31.84 c | 9.20 cd | 226.40 f | 1.78 d | 27.66 d | 4.03 f |
| **T4** | 22.50 b | 8.00 ab | 5.00 c | 39.08 a | 11.60 c | 331.00 d | 1.87 c | 22.86 e | 6.19 d |
| **T5** | 21.84 b | 8.00 ab | 5.00 bc | 29.26 d | 14.60 b | 423.80 b | 2.04 b | 33.97 b | 8.64 b |
| **T6** | 26.48 a | 9.20 a | 5.40 b | 35.34 b | 16.80 a | 463.80 a | 2.19 a | 31.36 c | 10.16 a |

Mean with same letters in each column are not significantly different at $p \leq 0.05$ according to LSD test

A gradation of diversity in tissues of both cultivated and wild rice was observed in the Shannon diversity index. Diversity indices of bacterial endophytes varied within plant parts as well as between cultivated and wild rice. High Shannon diversity index was recorded in roots (*H* = 2.718 and 1.946) of cultivated and wild rice, followed by in stem (*H* = 2.659 and 1.609). Diversity was least in leaf (*H* = 2.393 and 1.386) of both cultivated and wild rice. Simpson index (1-D) was highest in roots (0.930 and 0.857) with a richness of 16 different species occurring in cultivated rice and 7 species occurring in wild rice. The overall richness of endophytic bacteria as revealed in the Refraction curve indicated species richness in cultivated rice with maximum species richness in roots (**Fig. 2 & Table. 3**).

**Fig. 2:**
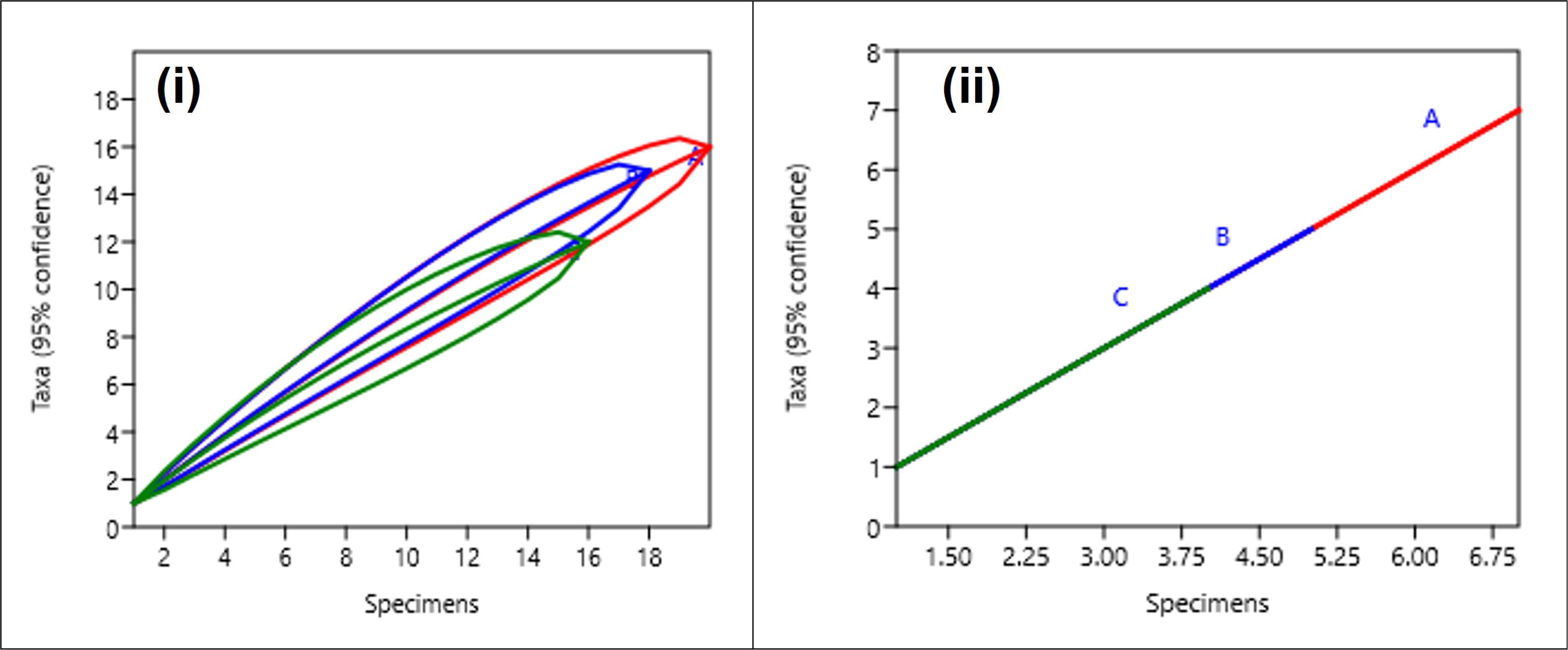
Rarefaction curve of bacterial endophytes: (i) cultivated and (ii) wild rice (A: root, B: stem and C: leaf).

Out of 54 endophytic bacteria isolated from different parts of cultivated rice, the isolates belonged to 11 different genera *viz. Bacillus, Brevibacillus, Lysenibacillus Microbacterium, Microbacteriaceae, Staphylococcus, Pantoea, Burkholderia, Acinetobacter, Pseudomonas*, and *Ralstonia*. While, *Bacillus, Stenotrophomonas, Microbacterium, Cellulosimicrobium, Proteus, Staphylococcus, Erwinia, Ochrobactrum*, and *Enterobacter* were the genera isolated from plant parts of wild rice (**Fig. 3(a)**). *Bacillus* sp., *B. pumilus, B. cereus, B. safensis, B. megaterium, Microbacterium* sp. were the most frequently occurring species found in stem, leaf, and root of *Oryza sativa. Bacillus megaterium* was found to be dominant species in the leaf while *Bacillus cereus* in the stem. *Bacillus cereus, Cellulosimicrobium cellulans*, and *Stenotrophomonas maltophilia* were found to be dominant species occurring in most plant parts of wild rice (**Fig. 3(b)**.

**Fig. 3:**
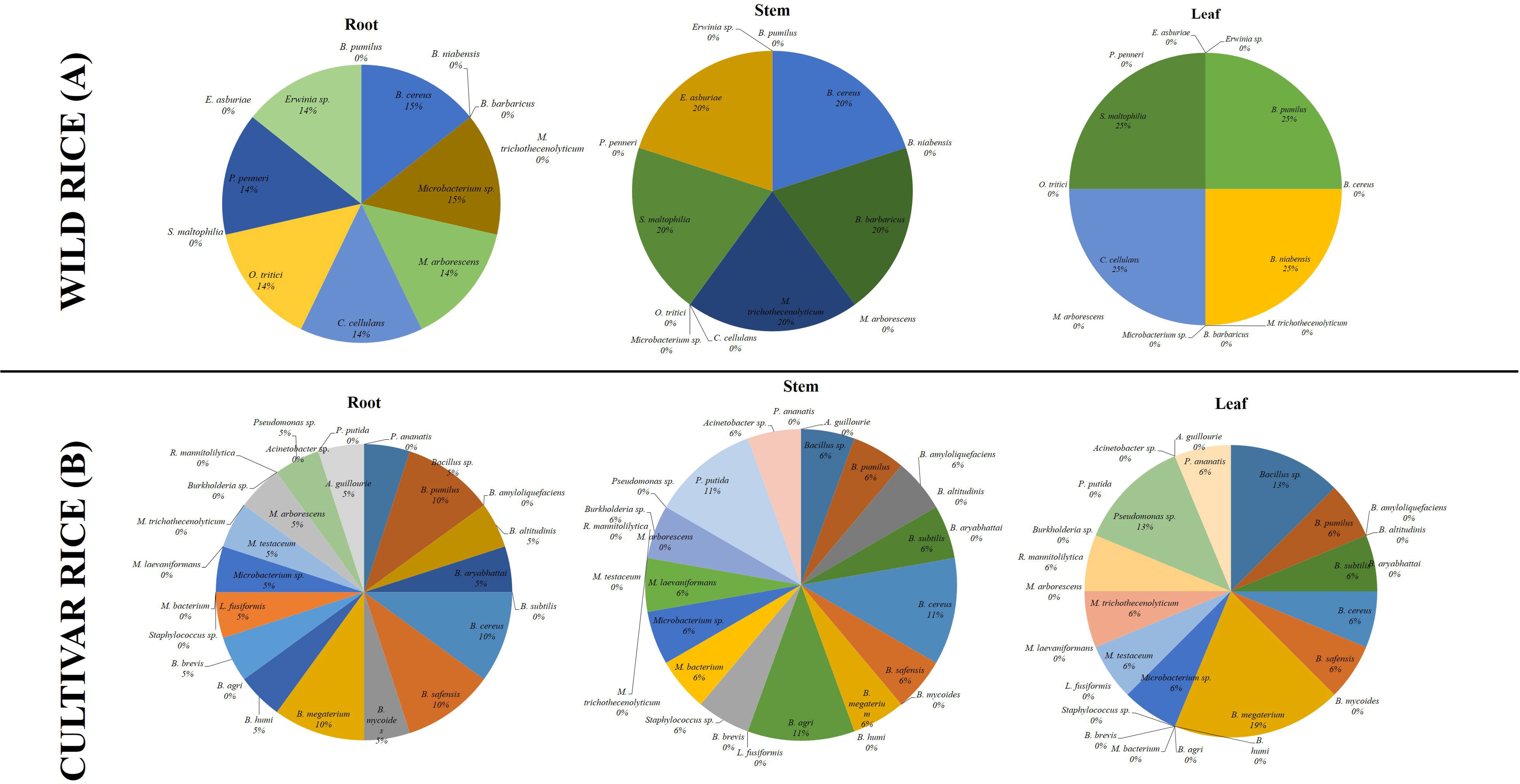
Diversity of bacterial endophytes isolated from different parts of (A) Wild rice and (B) Cultivar rice variety.

### Screening of isolates for plant growth promoting properties (PGP) Phytohormone Production Indole Acetic Acid (IAA)

The production of IAA is an important property of endophytic bacteria which aid in promoting plant growth. *In vitro* screening for IAA production revealed a substantial variation in the range of IAA (2.33 - 28.39 μg/ml) production among the 35 isolates that produced the phytohormone (**Fig. 4(a)**). The isolate, *Microbacteriaceae bacterium* RS01 11 produced significantly (*p* ≤ 0.05) higher amount of IAA (28.39 ± 1.33 μg/ml) when compared to other isolates.

**Fig. 4:**
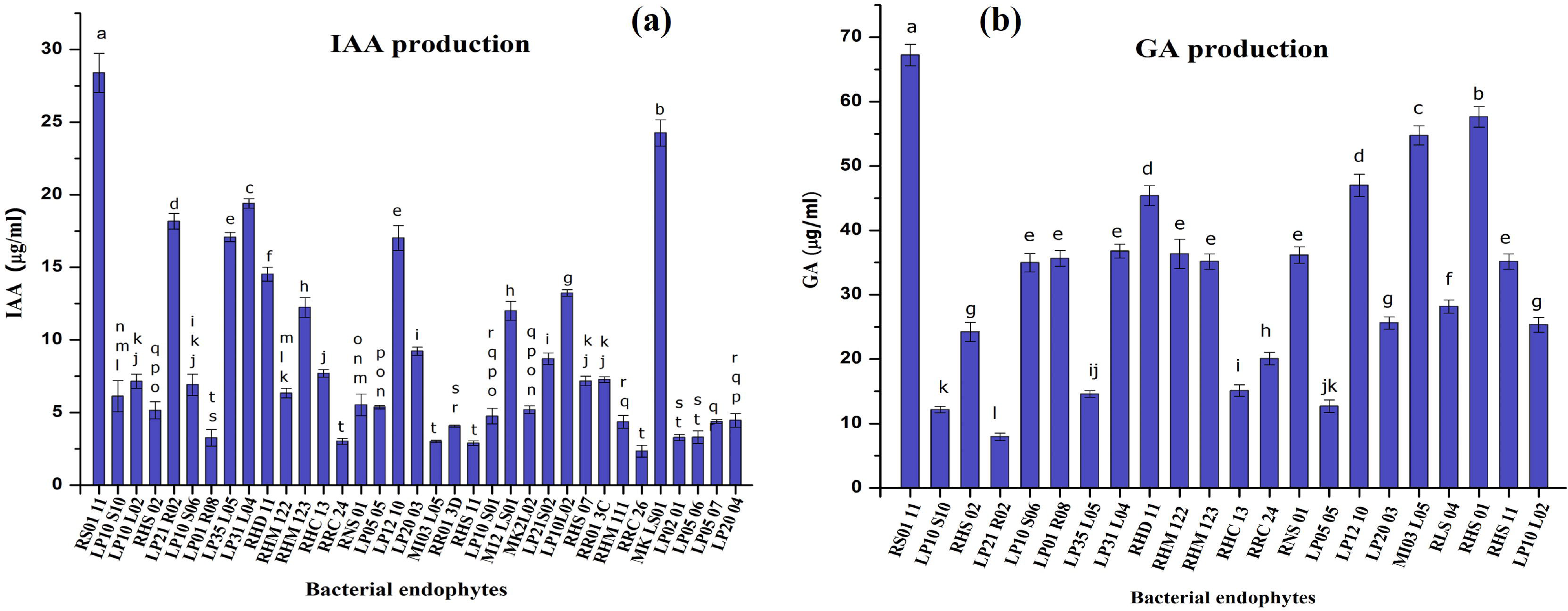
hytohormone production by the isolated bacteria (a) Indole acetic acid (IAA) (b) Gibberellic acid (Ga).

### Gibberellic Acid (GA) Production

Twenty four isolates showed the ability to produce gibberellic acid (GA) that ranged between 7.94 – 67.23 μg/ml (**Fig. 4(b)**). The isolate *Microbacteriaceae bacterium* RS01 11 also produced significantly (*p* ≤ 0.05) higher amount of gibberellic acid (67.23 ± 1.67 μg/ml) when compared to the isolate *Microbacterium testaceum* LP21 R02 that produced the least (7.94 ± 0.56 μg/ml).

### Mineral Solubilisation Phosphate Solubilization

Bacterial endophytes were screened for their ability to solubilize phosphate which revealed 35 isolates as potential phosphate solubilizer. Quantitative analysis of phosphate solubilizing abilities of these bacteria varied between 6.8 and 81.70 μg/ml (**Fig. 5(a)**). Phosphate solubilization activity was shown significantly (*p* ≤ 0.05) higher in *Bacillus subtilis* RHS 01 (81.70 ± 1.3 μg/ml) followed by *Brevibacillus agri* RS01 05 (60.75 ± 0.24 μg/ml).

**Fig. 5:**
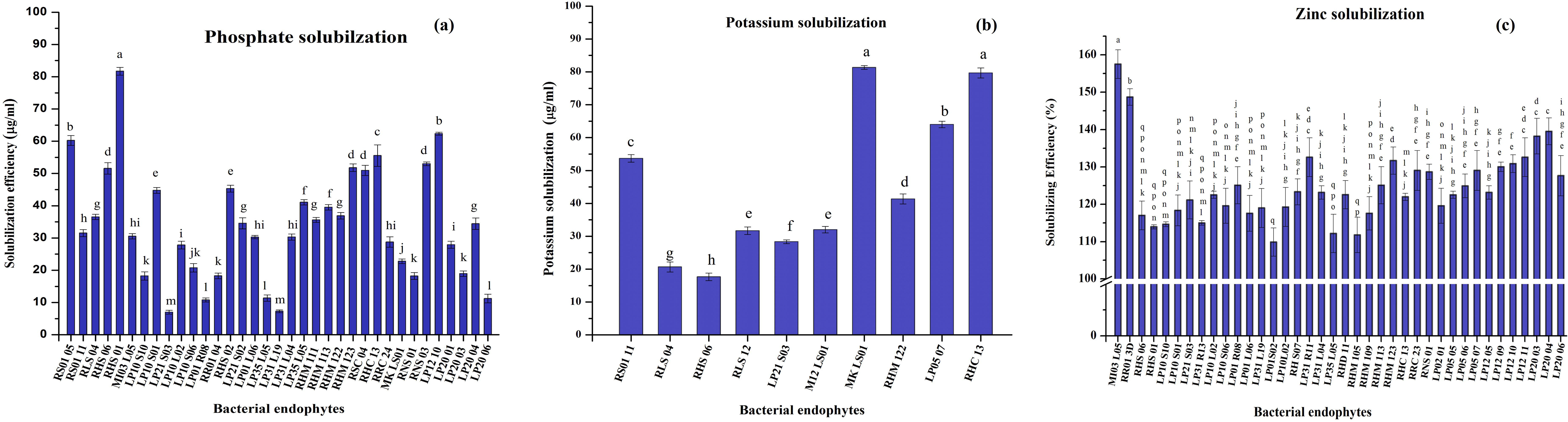
Mineral solubilization efficiency of the isolate endophytes (a) Phosphate (b) Potassium and (c) Zinc.

### Potassium Solubilization

Of the seventy isolates, ten isolates were found to be potential potassium solubilizers (**Fig. 5(b)**). The solubilizing efficiencies of the isolates ranged between 17.67 – 81.33 μg/ml. *Microbacterium testaceum* MK LS01 (81.33 ± 0.58 μg/ml) and *Bacillus cereus* RHC 13 (79.66 ± 1.67 μg/ml) had significantly (*p* ≤ 0.05) higher ability to solubilize potassium while *Bacillus pumilus* RHS 06 showed the least ability to solubilize potassium (17.67 ± 0.45 μg/ml).

### Zinc Solubilization

The bacterial isolates were inoculated in theTris minimal agar medium containing two different insoluble sources ZnO and ZnS of Zn at 0.1%. However, the endophytic isolates could solubilize only ZnO. The solubilization efficiency of the isolates was calculated by measuring the diameter of the colony growth and the solubilization zone. Zinc solubilizing efficiency of the isolates ranged between 110 % and 157.50 % (**Fig. 5(c)**). Zinc solubilization efficiency was found significantly (*p* ≤ 0.05) higher in *Microbacterium trichothecenolyticum* MI03 L05 (157.50 %) when compared to *Bacillus altitudinis* RR01 3D (148.72 %) and *Staphylococcus* sp. LP01S02 (109.89) with least solubilizing efficiency (**Fig 5(c)**).

### Siderophore production

For the initial detection of siderophore, bacterial endophytes were grown in modified Fiss minimal medium under low iron conditions. Fourteen bacterial endophytic isolates produced siderophore (Carson et al., 2000). This was further confirmed by CAS shuttle assay in which siderophore production was calculated in terms of percentage of siderophore units (**Fig. 6**). Siderophore units ranged between 7.06 - 64.80 %. *Bacillus barbaricus* LP20 05 produced significantly (*p* ≤ 0.05) higher siderophore units (64.8 %) when compared to *Bacillus megaterium* RLS 12 (7.06 %).

**Fig. 6:**
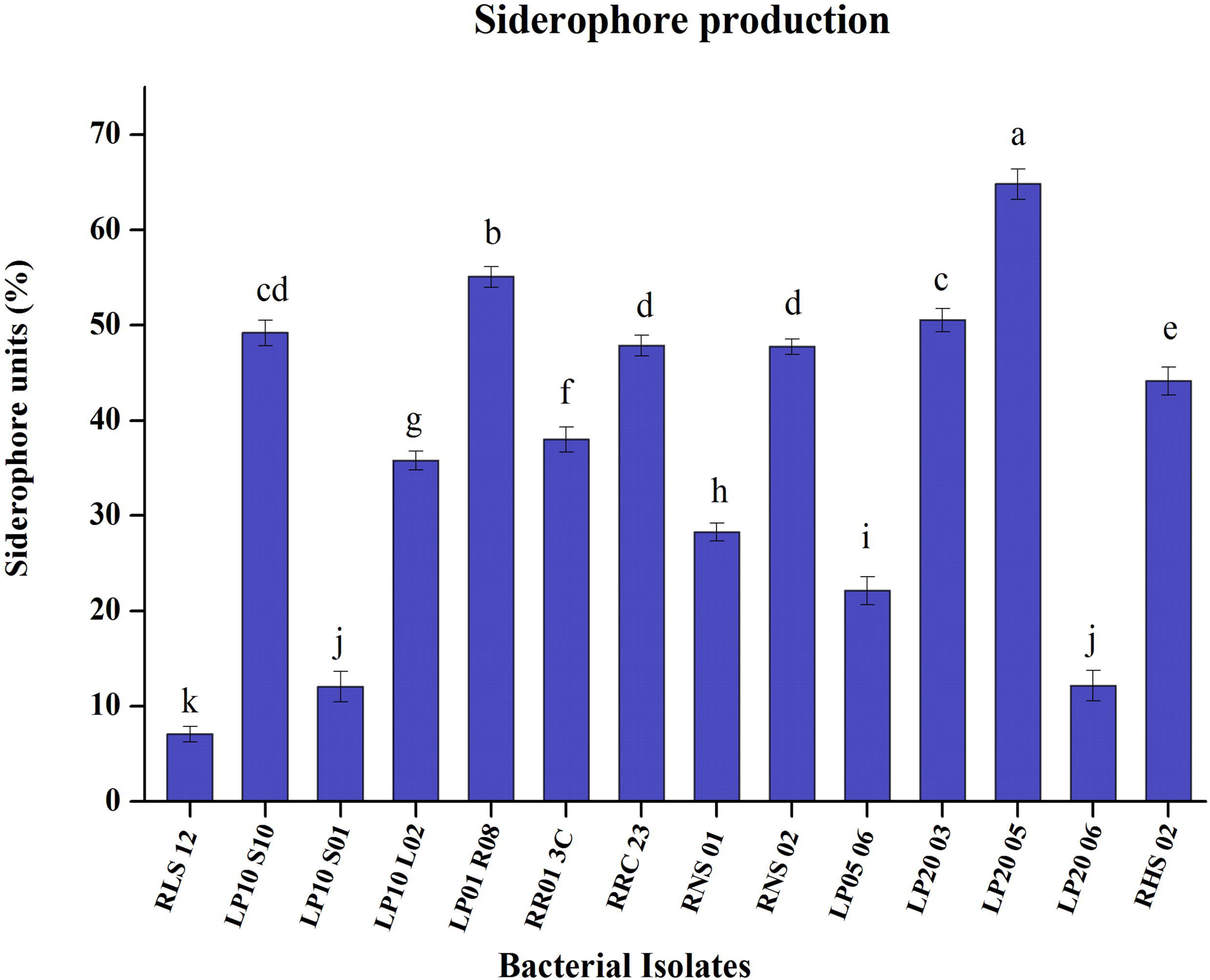
Siderophore production efficiency of the isolated endophytes.

### Efficacy of bio-inoculum on plant growth promotion under greenhouse conditions

Three endophytic strains namely, *Microbacteriaceae bacterium* RS01 11, *Bacillus subtilis* RHS 01 and *Microbacterium testaceum* MK LS01 were selected based on maximum PGP activity for their ability to promote plant growth in rice variety Dichang in pot culture experiment. The performance of the treatments in all combination was analyzed till harvest.

All the six treatments showed varying degree of growth as compared to the control (T0) treatment. The treatment T6 resulted in significantly (*p*≤0.05) higher shoot height, number of panicles per plant, the total number of seeds per plant, the weight of 100 seeds and yield per plant and with an average value of 26.48 cm, 16.80, 463.8, 2.19 gm and 10.16 gm respectively when compared to all other treatments. Length of flag leaf and dry biomass of the plant were highest in T2 and T6 treatment (**Table 4**). The pairwise comparison of treatments did not reveal any significance among the treatments for a number of tillers and number of leaves, although all treatments resulted significantly different from control T0 (**Fig. 7**).

**Fig. 7:**
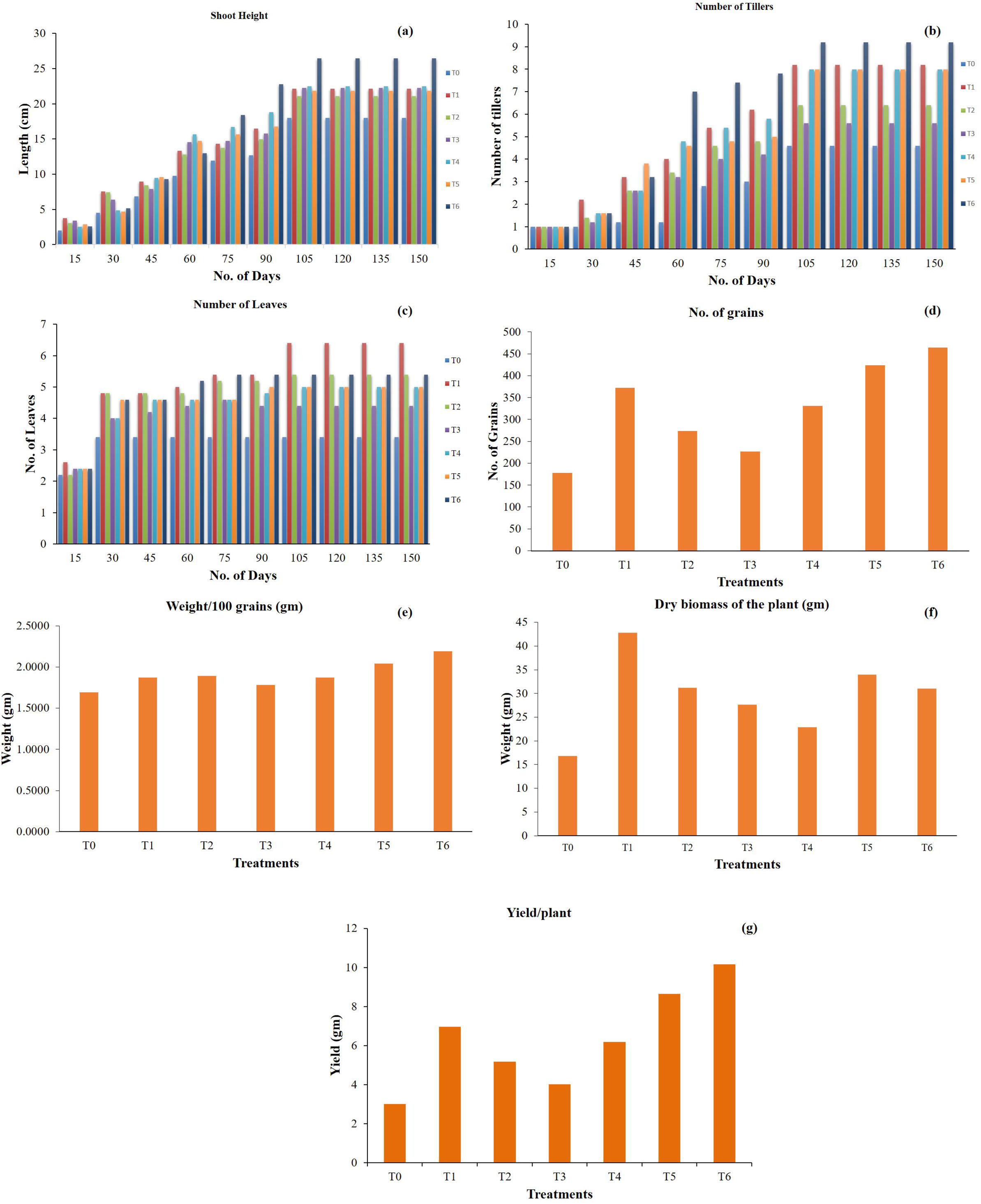
Pot culture experiment – evaluation of plant growth promoting efficiency of the isolates using rice as an test plant under controlled greenhouse environment. Parameters evaluated for the experiment, (a) Shoot Height of Rice; (b) Number of Tillers developed in Rice; (c) Number of Leaves; (d)No. of grains; (e) Wieght of 100 grains; (f) Dry biomass of the plant (gm); (g) Yield per plant.

## Discussion

Endophytes are the diverse group of endosymbiotic microorganisms which can directly or indirectly influence the growth and development of plants without causing any pathogenic effect on the host. Identification and characterization of diverse endophytic microorganism from different niches with potential phytostimulent activity can aid in improving sustainable agricultural practices. Morphological and biochemical characterization of the isolates revealed the dominance of gram-positive bacteria over the gram-negative bacterial endophytes. Molecular characterization of the isolates using 16S rRNA gene showed Firmicutes (57.1%) as the major colonizer of rice tissue particularly the *Bacillus* sp. Earlier reports suggested *Bacillus* as an efficient tissue colonizers in different plants including *Coffea arabica* L., sunflower, cotton, potato, strawberry, *Panaxnoto ginseng* and citrus plants (Araújo et al., 2001; Dias et al., 2009; Forchetti et al., 2007; Ma et al., 2013; Misaghi and Donndelinger, n.d.; Sessitsch et al., 2004; Vega et al., 2005). Production of a multilayered cell wall structure, formation of stress-resistant endospores and secretion of peptide antibiotics, peptide signal molecules, and extracellular enzymes are some of the physiological traits of *Bacillus* that enable them to survive in several different ecological niches (Lyngwi and Joshi, 2014). Other important phylum identified were and Actinobacteria (20%) and Proteobacteria (22.8%). Phylum Actinobacteria was represented by the family Microbacteriaceae under which *Microbacterium, Microbacteriaceae*, and *Cellulosimicrobium* were observed. Several species of *Microbacterium* was previously isolated from plants such as maize, rice and wheat (Conn and Franco, 2004; Elbeltagy et al., 2001; Rijavec et al., 2007). Proteobacterial sub-classification showed the dominance of Alphaproteobacteria, Betaproteobacteria, and Gammaproteobacteria. Pseudomonadaceae family with *Pseudomonas* as a major member of endophytes from proteobacteria accounted for 10 % of the total isolates. Genus *Pseudomonas* is a widely distributed plant-associated bacterium with reported activity of growth promotion in plants such as alfalfa (Gagné et al., 1987), clover (Sturz et al., 1997), potato (Reiter et al., 2002), and pea (Elvira-Recuenco and van Vuurde, 2000). Some minor groups such as Enterobacteriaceae, Moraxellaceae, Xanthomonadaceae, Burkholderiaceae were also observed from proteobacterial phylum.

Endophytic bacterial diversity was measured in terms of Shannon (*H*) and Simpson (1-D) diversity indices which indicated differences in cultivated and wild rice and species richness. Higher values of Shannon and Simpson indices are representative of more diverse communities. High indices were noted for roots of cultivated (*H* = 2.718, 1-D=0.930) and wild (*H*= 1.946, 1-D=0.857) rice. This could be explained on the basis that most endophytic bacteria are derived from the soil. The rhizosphere is the region for bacteria to reside and obtain nutrients (Raaijmakers et al., 2002). Bacteria residing in the rhizosphere might also have the potential to enter and colonize the plant roots. In fact, microbial population and their diversity in the rhizosphere is a major contributor for number and diversity of endophytes in a host plant (Hallmann and Berg, 2006). Some rhizoplane-colonizing bacteria can penetrate plant roots, and some strains may move to stem and leaves, with a lower bacterial density in comparison to root-colonizing populations (Compant et al., 2010). In the present study decrease of endophytic population was recorded from root onwards to the leaf through the stem. The reason maybe that most of the endophytes enter into the plant tissue through root and only a few can penetrate the xylem vessels through the casparian strip. The few microbes enter in to the xylem vessels slowly move towards the apical parts of the plant and hence the concentration of microbes decreases from root to stem and leaf (Gasser et al., 2011). In a study by Prakamhang *et al*. (2009) endophytic bacteria in rice were found in highest density in roots than other parts of the plant (Prakamhang et al., 2009).

All plants from cultivated to wild possess diverse endophytic microbiome. Such endophytes are of particular interest because they have high potential to produce different phytohormones and phytostimulatory compounds for promoting growth and yield. However, often these microorganisms are studied as a collective group and endophytes colonizing in different parts of a plant are rarely being analyzed. The study of microbial community in leaf, stem, and roots showed a difference in the microbiome of cultivated and wild rice. The microbial load in the wild rice was significantly less than the cultivated rice indicating the difference in their habitat throughout influence endophyte diversity. Wild rice is mainly found in the waterlogged marshy soils where abiotic stress is common, while cultivated varieties are traditionally grown with all the required nutrient supplements. Moreover, the cultivated rice seeds come from different locations which also influence upon the endophytic bacterial diversity.

The genus wise classification showed wild rice was mainly dominated by *Bacillus, Microbacterium, Cellulosimicrobium, Ochrobactrum, Pantoea, Enterobacteriaceae*, and *Erwinia*. However, cultivated verities showed a varied diversity of *Bacillus* along with *Microbacterium, Microbacteriaceae, Pantoea, Burkholderia, Acinetobacter, Pseudomonas, Acidovorax, Ralstonia, Staphylococcus, Lysenibacillus*, and *Brevibacillus*. Further annotation showed the variation is not only in the microbiome of cultivated and wild rice but there is a visible differentiation in the colonizing pattern of microorganism across the different parts of plants. Bacteria like *B. altitidinus, B. aryabhattai, B. mycoides, L. fusiformis*, and *A. guillouiae* are root-associated bacteria in cultivated rice while *O. tritci, P. penneri, Erwinia* sp., *Microbacterium* sp. were from wild rice. Interestingly *M. arborescens* is strictly root-associated bacteria in both the cultivated and wild rice. In significance, the colonization in the root is not random, like *M. arborescens* is beneficial to the root as they can produce high exopolysaccharide which helps in the soil aggregation and reports also suggest their involvement in iron-translocation in the rhizosphere. Microorganism mainly *B. mycoides* helps in nitrogen fixation, *B. altidins* can produce glucanase which helps the plant to inhibit the soil-borne pathogenic fungi, *B. aryabhattai* shows tolerance against nitrosative stress which protects the root cells from cellular damage. Stem specific diversity showed the dominance of *B. amyloliquefaciens, B. agri, Staphylococcus sp., M. bacterium, M laevaniformans, Burkholderia sp., P. putida* and *Acinetobacter sp*., in cultivated rice while *B. barbaricus, M. trichothecenolyticum*, and *E. asburiae* were dominant in the stem of wild rice. Leaf associated bacteria was not as diverse as other parts, *P. ananatis, R. mannitolilytica, M. trichothecenolyticum* was found in cultivated rice while *B. pumilus*, and *B. niabensis*, was found in wild rice. Bacteria found on the leaf of cultivated varieties were a mostly opportunistic human pathogen, which suggests the human intervention in the farmland, whereas *B. pumilus* and *B. niabensis* are native plant associate encouraging the less anthropogenic activities. Other identified bacteria from cultivated and wild rice like *Bacillus sp., B. subtilis, B. cereus, B. safensis, B. megaterium, Pseudomonas sp., B. cereus, C. cellulans, M. testaceum* and *S. maltophilia* showed abundance in all the parts with exceptions. *Bacillus pumilus*, which is dominant in all three parts of the plant in cultivated rice but only leaf specific in wild rice.

Endophytes can directly or indirectly influence on growth and development of plants without causing any pathogenic effect on the host. The mutual interaction is habitually facilitated by a number of metabolites linked with impelling mechanisms *viz*. mobilizations and uptake of nutrients and production or co-regulation of phytohormones. Phytohormone production by endophytes is perhaps the best-studied mechanism of plant growth promotion (Long et al., 2008). Indole-3-acetic acid (IAA), the most common auxin found in the plant (Barazani and Friedman, 1999) regulates various aspects of plant growth and development (Bulgarelli et al., 2013). It acts as a regulator of numerous biological processes such as cell division and elongation, tissue differentiation, apical dominance, and responses to light, gravity, and pathogens (Aloni et al., 2006). Our study revealed that *Microbacteriaceae bacterium* RS011 produced a significant amount of Indole-Acetic-Acid (28.39±1.33 μg/ml) followed by *Microbacterium testaceum* MK LS01 (24.25±0.90 μg/ml). *Bacillus* sp. isolated from rice had been reported earlier for their IAA producing activity by Phetcharat and Duangpaeng (2012) (Phetcharat and Duangpaeng, 2012). Gibberellic acid (GA), a class of major phytohormone responsible for seed germination and mobilization of food-substances for the growth of a new cell, was found to be maximum in *Microbacteriaceae bacterium* RS01 11 (67.23 ± 1.67 μg/ml) followed by *Bacillus subtilis* RHS 01 (57.63 ± 1.57). Gibberellic acid has also been reported to be synthesized by several bacterial species including *Acinetobacter, Azospirillum brasilense, Agrobacterium, Arthrobacter, A. lipoferum, Azotobacter, Bacillus, Bradyrhizobium japonicum, Flavobacterium, Micrococcus, Clostridium, Pseudomonas, Rhizobium* and *Xanthomonas* (Gutierrez-Manero et al., 2001). Endophytic *Pseudomonas* and *Bacillus* isolates of tropical legume crops were also reported to secrete GA (Maheswari and Komalavalli, 2013). To the best of our knowledge, this is the first report on rice endophytic *Microbacteriaceae bacterium* with the ability to secrete substantial amount of IAA and GA.

Plant growth and yield are essentially dependent on the availability of minerals which they directly or indirectly acquire from the soil. Soil constitutes 0.5% phosphorus, mostly in the form of insoluble mineral complexes which plants cannot directly absorb (Rengel and Marschner, 2005), only 0.1 % of the total P exists in a soluble form available for plant uptake (Sharma et al., 2013). Phosphate-solubilising bacteria (PSB) are able to solubilise bound phosphorous from organic or inorganic molecules, by secretion of organic acids such as phytic acid, formic acid, acetic acid, lactic acid and by producing enzymes such as phosphatases or C-P lyases (Chung et al., 2005; Kim et al., 1998). Characterization of PSB showed *Bacillus subtilis* RHS 01 as a potent phosphate solubilizer (81.70±1.3 μg/m) followed by *Brevibacillus agri* RS01 05 (60.75±0.24 μg/ml). These endophytic bacterial isolates were able to solubilize organic or inorganic form of phosphate suggesting that they could play a role in resource mobilization in nutrient-poor habitat. The previous study by Dias *et al*., (2009) confirmed the efficiency of endophytic bacteria such as *Bacillus subtilis* and *B. megaterium* isolated from strawberry in solubilization of tricalcium phosphate (Dias et al., 2009).

Potassium is a major nutrient for plant growth and development, which provides an ionic environment for metabolic process in the cytosol and as such function as a growth regulator. *Microbacteriaceae testaceum* MK LS01 (81.33 ± 0.58 μg/ml) and *Bacillus cereus* RHC 13 (79.66 ± 1.67 μg/ml) solubilized the highest amount of potassium in the study. *Bacillus cereus* isolated from soil is reported to solubilize potassium (Diep and Hieu, 2013). Yuan *et al*., 2015 isolated 14 species from 10 genera of potassium-solubilizing endophytic bacteria from moso bamboo which mainly consist of *Alcaligenes* sp., *Enterobacter* sp. and *Bacillus* sp. Other genera such as *Burkholderia* sp., *Paenibacillus* sp., and *Acidothiobacillus* sp. were reported to be potassium solubilizing biofertilizer (Nair and Padmavathy, 2014). *Microbacterium foliorum*, isolated from tobacco rhizosphere has the ability to solubilize potassium (Zhang and Kong, 2014). However, this is the first report of potassium solubilization property of endophyte *Microbacteriaceae testaceum*, isolated from rice.

Zinc, though a micronutrient is one of the essential minerals for chlorophyll synthesis. Zinc solubilizing microorganisms have the ability to dissolve the immobilized zinc *viz*. zinc phosphate, zinc oxide and zinc carbonate in considerable quantity (Saravanan et al., 2007). In the present study, *Microbacterium trichothecenolyticum* MI03L05 and *Bacillus altitudinis* RR03D showed a significant amount of zinc solubilization with an efficiency of 157.50 % and 148.64 % respectively. The formation of halo zones by the microorganisms is due to the movement of acidity corresponded with the solubilization of the metal compound (Fasim et al., 2002). Other bacterial genera viz. *Acinetobacter, Bacillus, Gluconacetobacter, Pseudomonas, Thiobacillus thioxidans, Thiobacillus ferroxidans*, and facultative thermophiliciron oxidizers have also been reported as zinc solubilizers (Saravanan et al., 2007). Endophytic *Bacillus* sp. and *Pseudomonas* sp. isolated from soybean were also reported to solubilize zinc (Ramesh et al., 2014). But there are no previous reports on zinc solubilization by *Microbacterium trichothecenolyticum* which is supposed to be the first of its kind.

Production of siderophore for iron chelation is an important trait in endophytic bacteria. Despite being the most available element in the earth crust, the bioavailability of iron is very limited due to the low solubility of Fe^+3^ ion and siderophores, perhaps, are the strongest binding agent of Fe^+3^. A number of plant species can directly absorb the bacterial Fe^3+^ siderophores complexes (Beneduzi et al., 2012). *Bacillus barbaricus* LP20 05 (64.8 %) was found to exhibit highest siderophore production activity, followed by *Lysenibacillus fusiformis* LP01R08 (55.05%). Endophytic bacteria *Bacillus barbaricus* isolated from *Zingeber officinale* was reported to produce siderophore (GINTING et al., 2013). Siderophore production is also shown by rhizospheric bacteria *Bacillus barbaricus* (21%) and *Pseudomonas fluorescens* (76%) (Gupta and Gopal, 2008).

Increase application of PGPB can be seen in sustainable agricultural practices for growth enhancement and increased crop yield (Kloepper et al., 1991). In the present study, the pot culture evaluation revealed the T6 treatment (Soil + ½ NPK + ½ Vermicompost + ½ Bioinoculum) as effective in promoting plant growth in terms of shoot heights, number of tillers, number of panicles, the total number of seeds per plant, weight of 100 grains and yield per plant. The T6 treatment resulted in significantly (*p* ≤ 0.05) higher increase on growth and yield parameters when compared to control T0 and T1 (NPK). The delay in acclimatization and colonization of the microorganisms in soil and rhizosphere may initially take time to show the benefit, however once established the PGPB enhanced plant growth and yield. Endophytic PGPB are good candidates to be used as inoculant as they can colonize roots and create a favourable environment for development and yield (Bacon and Hinton, 2007). The mechanisms by which bacteria can influence plant growth differ among species and isolates, usually there is no single mechanism for promoting plant growth (Souza et al., 2015) hence mechanisms that stimulated plant growth could be explained by combined effects of all the PGPR properties like phytohormone (IAA and GA) production and mineral (phosphate, potassium, and zinc) solubilisation of each isolate present in the bio-inoculum (Bashan et al., 2004; Glick, 1995). *Microbacteriaceae bacterium* RS01 11 had phytohormone biosynthesis capacity which might have stimulated the process of plant growth. The secretion of IAA might have aided in improving root development while GA in promoting shoot growth. *Bacillus subtilis* RHS 01 and *Microbacterium testaceum* MK LS01 were able efficient phosphate and potassium solubilizers respectively. *Bacillus subtilis* RHS 01 and *Microbacterium testaceum* MK LS01 had good zinc solubilising activity as well that might have helped the process of growth enhancement. Earlier greenhouse and pot culture studies with the endophytic rhizobial inoculum indicated the significant increase in N, P and K uptake in rice plants and led to increased biomass and yield of rice plants (Biswas et al., 2000). Similar kind of experiment was conducted by Ashrafuzzaman et al., (2009) to determine the efficiency of plant growth-promoting rhizobacteria (PGPR) for the enhancement of rice growth and found that the use of PGPR isolates PGB4, PGG2, and PGT3 as inoculant biofertilizers were beneficial for rice cultivation as they enhanced the growth of rice.

## Conclusion

The study generated a baseline data on the endophytic bacterial diversity of cultivated and wild rice in Assam through a culture dependent method. The endophytes *Microbacteriaceae bacterium*, *Microbacterium testaceum* and *Bacillus subtilis* exhibiting multiple plant growth promoting activity with high efficiency can form a prospective consortium which can further be employed as a source of bio-fertilizer for enhancement of plant growth and development. Current research also suggests that the inoculation of crops with bioinoculum of endophytic bacteria has the potential to reduce application rates of chemical fertilizer to half the recommended dose.

## Acknowledgment

Authors are grateful to the funding organization Department of Biotechnology (DBT), Ministry of Science and Technology, Govt. of India and Department of Agricultural Biotechnology, Assam Agricultural University, Jorhat, Assam for carrying out the research.

